# Association between loneliness and hippocampal responses to dynamic social stimuli in psychotic disorders

**DOI:** 10.1101/2024.10.31.621352

**Authors:** Faye McKenna, Louis N. Vinke, Mona Avanaki, Daphne J. Holt

**Affiliations:** Department of Psychiatry, Massachusetts General Hospital, Boston, MA, USA; Harvard Medical School, Boston, MA, USA; Athinoula A. Martinos Center for Biomedical Imaging, Massachusetts General Hospital, Boston, MA, USA

**Keywords:** loneliness, isolation, functional MRI, trust, psychotic disorders, hippocampus, basal ganglia

## Abstract

**Background:** The incidence of loneliness has increased over the past several decades worldwide and is particularly common among people with serious mental illnesses. However, this public health problem has been difficult to address, in part because the neurocognitive mechanisms underlying loneliness are poorly understood.

**Methods:** To investigate these mechanisms, a functional magnetic resonance imaging (fMRI) study was conducted which accounted for known cognitive biases associated with loneliness. Participants with (n = 40) and without (n = 60) psychotic disorders (PD) viewed images of faces that appeared to approach or withdraw from the participants, while fMRI data were collected. Following the scanning, participants rated the trustworthiness of the faces, and these ratings were included as weights in the fMRI analyses. Neural responses to approaching versus withdrawing faces were measured, and whole brain regression analyses, with loneliness as the regressor, were performed.

**Results:** In the PD and full samples, a higher level of loneliness was significantly associated with greater responses of the hippocampus and areas of the basal ganglia to withdrawing (versus approaching) face stimuli. Moreover, the effects in the hippocampus, but not the basal ganglia, remained significant after controlling for potential confounds such as social activity levels, depression and social anhedonia. Lastly, in a subset of the sample (n = 66), greater hippocampal responses to withdrawing faces predicted greater loneliness one year later.

**Conclusions:** Heightened responses of the hippocampus to withdrawing faces may represent a candidate objective marker of loneliness that could be modified by interventions targeting loneliness.

## INTRODUCTION

Loneliness has been defined as a perceived deficit in social connection (a discrepancy between desired and actual levels of social connection), which can be distinguished from social isolation, an objective deficit in social activity (1). In recent years, public health experts have called attention to a worsening **“**epidemic” of loneliness and social isolation in the general population that has been ongoing for decades and then exacerbated by the COVID-19 pandemic (2,3). In addition, it has been well-established that loneliness and isolation are common among those suffering from psychiatric illnesses, including psychotic disorders, affecting up to 80% of people with these conditions (4). This widespread societal problem has broad public health implications, since numerous epidemiological studies have linked loneliness and isolation to poor cardiometabolic health and early mortality (5). However, little is known about the psychological and neurobiological underpinnings of these associations and how they might vary across different populations.

Some clues about the psychological mechanisms of loneliness have emerged from studies showing that certain cognitive and behavioral biases are associated with loneliness, including a bias towards social mistrust (6) and a related tendency to prefer a greater amount of physical space (“personal space”) from others (7,8). Specifically, loneliness has been linked to a sensitivity to social rejection and a related mistrust of others’ intentions (6), which can paradoxically lead to social withdrawal or avoidance of social activity (and thus greater isolation and loneliness).

In the current study, we employed a fMRI paradigm and analytic approach that accounted for these biases. During scanning, images of human faces were presented that appeared to move towards and away from participants, crossing participants’ personal space boundaries (9–12). Since an enlarged or more rigid personal space may represent a behavioral marker of the cognitive biases associated with loneliness (7,8), we hypothesized that this fMRI paradigm engages some of the neural circuitry involved in the experience of loneliness. Following scanning, participants also rated the trustworthiness of the face stimuli they had viewed, and these ratings were incorporated in the fMRI analyses, based on the hypothesis that neural responses to the faces rated as least trustworthy are most relevant to the experience of loneliness. Thus, using this approach, we sought to identify neural responses associated with loneliness in individuals with and without psychotic disorders.

The findings of prior neuroimaging studies of loneliness have been varied, with reports of associations between loneliness and responses or volumes of medial temporal lobe regions (6,13–21), basal ganglia (6,14–16,20,22) or the hypothalamus (22). For example, several studies have identified associations between the size (18–21,23) or functioning (13,14,16,17) of the hippocampus and loneliness. In addition, studies conducted in rodents have found evidence for a central role of the amygdala (24) and hypothalamus (24) in responses to social isolation, and studies in humans have also shown that the size (25) and connectivity (25) of the amygdala are linked to social activity levels. Thus, in the current study we tested the hypothesis that stronger associations between loneliness and responses of these previously implicated brain areas would be detected using an fMRI paradigm that appears to violate personal space boundaries, triggering a sensitivity to social threat. Additionally, because loneliness tends to correlate with social isolation as well as depression and social anhedonia, we conducted secondary analyses that controlled for these variables, with the goal of isolating specific associations with loneliness. Lastly, in a subset of individuals with longitudinal data, we tested if the responses of these brain areas predicted changes in loneliness over a one-year period.

## METHODS

### Participants

#### Recruitment

Sixty healthy control subjects and forty subjects with a history of a diagnosis of a psychotic disorder (PD) were recruited and enrolled in the current study (see Table 1 for participant characteristics). Of the forty PD participants enrolled, three were later excluded due to having incomplete fMRI data (see Supplementary Methods for data quality assessment exclusion criteria). A research domain criteria (RDoC) approach (26) was adopted for the design of this study, i.e., the PD category was defined broadly, including individuals with diagnoses of schizophrenia (*n* = 17), schizoaffective disorder (n=9), and bipolar disorder with a history of psychosis (*n* = 11). In addition, analyses were conducted both within each group and in the full sample, in order to fully test our dimensional hypotheses regarding the neural mechanisms of loneliness.

**Table 1.** Demographic and clinical characteristics of the sample.

| Measure | HC (n = 60)<br>Mean (SD) | PD (n = 37)<br>Mean (SD) | p - value | Test statistic |
| --- | --- | --- | --- | --- |
| Gender (% female) | 35.0 | 29.7 | 0.592 | $\chi^2(1, N=97) = 0.288$ |
| Age (yrs) | 26.12 (5.53) | 27.81 (6.24) | 0.166 | $t(95) = -1.395$ |
| Mean parental education (yrs) | 15.58 (3.33) | 15.78 (2.86) | 0.762 | $t(95) = -0.304$ |
| IQ<sup>a</sup> | 109.92 (7.44) | 106.71 (9.75) | 0.072 | $t(95) = 1.822$ |
| Task performance<sup>b</sup> | 80.7% (8.0%) | 78.3% (10.2%) | 0.204 | $t(95) = 1.280$ |
| Trustworthiness rating<sup>c</sup> | 4.36 (0.17) | 3.98 (0.72) | 0.051 | $t(95) = -1.973$ |
| Personal space size<sup>d</sup> | 79.40 (36.1) | 107.25 (50.6) | 0.002* | $t(95) = -3.155$ |
| Loneliness at baseline<sup>e</sup> | 29.94 (10.86) | 39.61 (11.84) | < 0.001* | $t(95) = -4.175$ |
| Loneliness at follow-up<sup>f</sup> | 32.41 (12.02) | 39.60 (10.42) | 0.018* | $t(60) = -2.435$ |
| Loneliness change<sup>g</sup> | 1.76 (7.32) | 0.56 (9.35) | 0.575 | $t(60) = 0.564$ |
| Social activity levels<sup>h</sup> | 18.48 (7.23) | 15.58 (10.45) | 0.002* | $t(95) = 3.127$ |
| Depression<sup>i</sup> | 2.94 (5.27) | 7.84 (8.85) | < 0.001* | $t(95) = -3.892$ |
| Social anhedonia<sup>j</sup> | 7.89 (7.30) | 10.65 (7.78) | 0.089 | $t(95) = -1.719$ |
| Symptoms of schizophrenia<sup>k</sup> | — | 67.14 (17.8) | — | — |
| Illness duration (yrs) | — | 7.49 (6.03) | — | — |
| CPZ equivalents (mg) | — | 253.93 (352.83) | — | — |

Participants were recruited via advertisements in online community forums and postings on research portals (Rally; Clinical trials https://rally.massgeneralbrigham.org/). PD participants were recruited either through online advertisements or the MGH Psychosis Clinical and Research Program (PCRP). DSM-V diagnoses of all participants were evaluated by trained research staff using the Mini International Neuropsychiatric Interview (27).

In accordance with the Declaration of Helsinki, written informed consent was obtained from all participants before the beginning of the study. All study procedures were approved by the Massachusetts General Brigham Institutional Review Board. See the Supplementary Methods for additional inclusion/exclusion criteria.

#### Clinical assessment and loneliness and social isolation measures

In all participants, loneliness was assessed using the UCLA Loneliness Scale (28), which is a well-validated, self-report measure of loneliness that has been used in many populations including psychotic disorders (29,30). Social activity levels were measured using the Social Network Index (SNI), which assesses the number of social contacts that the participant had during the previous two weeks (31). Also, depression (32), social anhedonia (33) and the symptoms of schizophrenia (in the PD group only) were measured using well-validated instruments. In addition, the identical assessments were also administered one year following the baseline time point (collected in 35 HC and 31 PD participants who were willing to participate in the follow-up assessment).

#### Functional MRI task (the “Looming” paradigm)

At baseline only, participants underwent a MRI scan session which included a well-validated fMRI paradigm (9–12) during which participants view images of human faces (8 female and 8 male -appearing digital face images, each with a neutral facial expression, half with eyes open and half with eyes closed, created using https://facegen.com), which either increase in size (appear to be approaching the participant) or decrease in size (appear to be withdrawing from the participant) over the course of each 16 second trial. Each functional run included a total of 16 trials, with 16 second blank (neutral gray) fixation blocks presented at the beginning and end of each functional run. Face gender and looming trial type were randomized and counter-balanced within each functional run.

#### Attentional task during scanning

While maintaining central fixation throughout each functional run, participants were instructed to covertly attend to other areas of the screen and report whenever they detected the appearance of a dot at a random location on the screen. During each 16-second looming trial, a dot appeared 3 times, with the duration between subsequent dot presentations randomly varying between 3.5 s and 4.5 s. Additionally, the duration of each dot presentation varied randomly between 350 ms and 1,500 ms, with the dot size scaled with eccentricity (i.e., larger diameter when presented further from central fixation). Participants responded using their right index finger to press a key on the response box provided.

#### Face ratings and personal space measurement

Following the scan session, each participant was shown the face stimuli that they had viewed during the scanning, one at a time, in a randomized order. Each of the 16 faces was shown four times. During the viewing of each face, participants rated (self-paced) the trustworthiness of the face (among other ratings) on a scale of 0-10, with 0 representing the least, and 10 the most, trustworthy. In addition, personal space size was measured on the same day as the scan session using the Stop Distance Paradigm, which is a validated, reliable procedure for measuring personal space preferences (see Supplementary Methods for details) (34).

### MRI Data Analysis

All MRI data analyses were conducted using FreeSurfer v7.4.1. See the Supplementary Methods for the standardized structural and functional MRI data preprocessing procedures and the data quality assurance methods used.

#### Regions-of-interest (ROI) definition

The Looming paradigm consistently engages a network of cortical and subcortical regions (9–12), which were defined in this study using 1) previously collected data that were acquired with this paradigm or 2) anatomical criteria. An independent fMRI dataset collected with the Looming paradigm (in a non-clinical sample of young adults (n = 130)) was used to independently define the cortical regions-of-interest (ROIs) (11). These *a priori* cortical ROIs included bilateral parietal and ventral premotor cortical regions (see Supplementary Methods). Activation in these cortical ROIs was assessed as a positive control, to establish that the task engaged the expected cortical regions (9–12). The hypothesis-testing analyses focused on *a priori* regions of subcortical networks that have been implicated in prior studies of the neural correlates of loneliness: the medial temporal lobe (the amygdala and hippocampus) (13–21,23), the basal ganglia (the caudate nucleus, putamen, nucleus accumbens, pallidum, and thalamus) and the basal forebrain, which includes the hypothalamus (6,14–16,20,22). These eight subcortical ROIs were defined using the automatic subcortical segmentation algorithm of Freesurfer (https://surfer.nmr.mgh.harvard.edu/).

#### Functional MRI data analyses

The first-level analysis used a univariate general linear model (GLM) fit to the event-related BOLD time-series data, for all runs passing quality assurance steps. The GLM included a canonical SPM hemodynamic response function, head motion (6 parameters) and scanner drift variables as nuisance regressors and excluded any time-points identified as outliers (see Supplementary Methods for additional details). The GLM contrast effect size (CES) maps were generated when contrasting responses of all approaching face trials versus all withdrawing face trials (i.e., Approach vs. Withdrawal contrast) using each participant’s individual trustworthy rating of each face as run-specific parametric regressors (e.g. a model comprised of the Approach vs Withdrawal contrast together with the trustworthy ratings for each subject) (35).

Both cortical and subcortical group-level analyses were performed to identify regions where the group-wise contrast effect size (CES) maps were significantly different from zero; significance maps were thresholded at *p* < 0.05, and false discovery rate (FDR) permutation-testing (cluster-wise *p*-value < 0.05) was applied for whole brain correction, for the cortical surface and subcortical space separately. Between-group differences were initially examined in the *a priori* ROIs using whole-brain voxel-wise comparisons conducted at a liberal *p* < 0.05 uncorrected threshold in order to facilitate comparisons to prior studies (36). These analyses were followed by analyses which 1) controlled for potential confounds and 2) used a whole-brain *p* < 0.05 cluster-wise FDR-corrected threshold.

#### Whole brain regressions using loneliness as a regressor

Regression analyses were conducted using a GLM voxel-wise technique utilizing the (CES) maps (Approach vs Withdrawal contrast X trustworthy regression coefficient) to measure the relationship between Looming activation and loneliness in the HC, PD, and full samples. Additional second-level GLM regression analyses, conducted in the full sample only, included 1) measures of potential confounds as covariates, and 2) change in loneliness from baseline to the one-year follow-up time point.

## RESULTS

### Loneliness, social activity levels, and clinical measures

Levels of self-reported loneliness were significantly greater on average in the PD group than in the HC group (*t*(97) = -4.175, *p* < 0.001; Table 1), and mean social activity levels were significantly lower in the PD, compared to the HC, group (*t*(97) = 3.175, *p* = 0.002). At the one-year follow-up timepoint, loneliness remained significantly greater in the PD, compared to the HC, group (*t*(60) = -2.435, *p* = 0.018), but the change in loneliness from baseline to one year was not significantly different between groups (*t*(60) = 0.560, *p* = 0.575) (Table 1).

Also, at baseline, loneliness was significantly correlated with both depression (all *r* > 0.712; all *p* < 0.001) and social anhedonia in each group and the full sample (all *r* > 0.630; all *p* < 0.001; Table S1). In addition, loneliness and social activity levels were negatively correlated with each other in the full sample (*r* = -0.323, *p* = 0.001; Table S1), but not in the HC and PD groups separately.

### Behavioral tasks

There were no significant differences between the HC and PD groups in performance on the attentional task during the scanning, and in the trustworthy ratings of the face stimuli viewed during the Looming task (Table 1, Figure S1).

### Neural responses to looming faces

#### Within-group effects

There were significant looming effects (Approach > Withdrawal) in all cortical ROIs in both groups and the full sample (Table S2 and Figure S2). With respect to subcortical areas, in the HC group, there were significant looming effects within the bilateral basal forebrain and right thalamus, left hippocampus, left caudate and left putamen (Table S2 and Figure 1), whereas in the PD group, significant effects were found in the left basal forebrain, bilateral thalamus, bilateral hippocampus and left amygdala (Table S2 and Figure 1).

**Figure 1.**
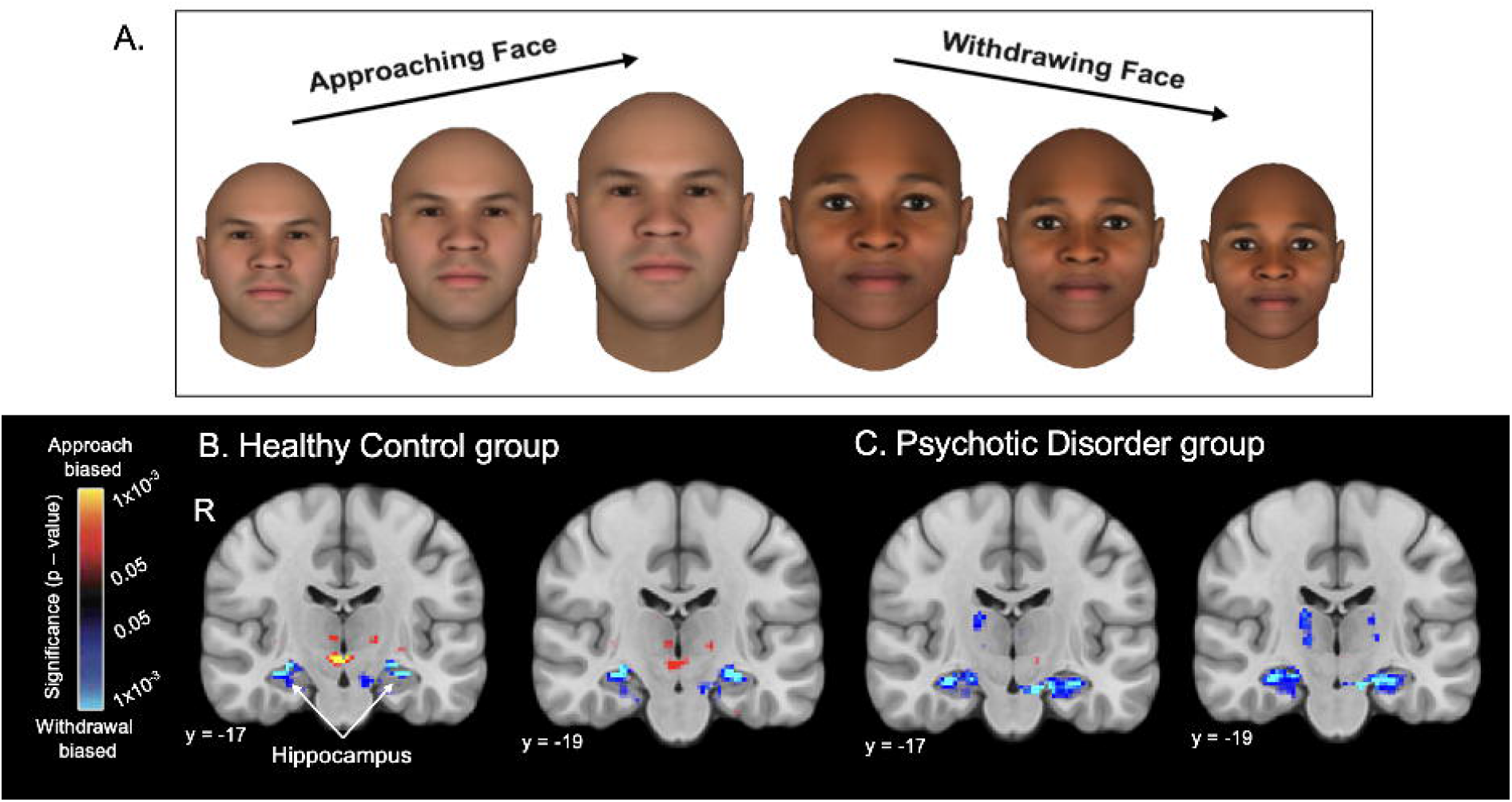
The Looming task stimuli and activation within the healthy control and psychotic disorder groups. A. Examples of the Looming task face stimuli are shown. During a trial, one unique face stimulus either increases (Approach condition) or decreases (Withdrawal condition) in size over the course of a 16 second block. The face displayed on the left (approaching, three images) was rated the least trustworthy face on average, while the face shown on the right (withdrawing, three images) was rated the most trustworthy face on average, across the full sample (n = 97). B. The functional contrast of Approach > Withdrawal was computed to assess neural responses to the Looming task. Subcortical voxel-wise activation (Approach > Withdrawal) maps are shown for the Healthy Control (n = 60) and Psychotic Disorder (n = 37) groups, with a significance map (display threshold: *p* < 0.05, uncorrected) overlaid on the MNI common space brain template, showing significant Withdrawal > Approach responses of the right and left hippocampus in two coronal slices (y = -17 and y = -19) in the two groups. Approach biased = Approach > Withdrawal; Withdrawal biased = Withdrawal > Approach; R, right hemisphere.

#### Between-group comparisons

At a liberal threshold (*p* < .05 uncorrected), some significant differences between the two group were evident in the bilateral basal forebrain, bilateral hippocampus, left amygdala, bilateral caudate and left putamen (Table S3 and Figure S3), with greater responses in these regions to Approaching > Withdrawing faces in the HC compared to the PD group (primarily due to elevated responses to the Withdrawing > Approaching faces in the PD group). These differences were not significant, however, after controlling for between-group differences in loneliness or social activity levels (Figure S3). Also, there were no significant differences between the two groups in activation in any of the ROIs at a *p* < 0.05 cluster-wise FDR-corrected threshold.

### Neural correlates of loneliness

#### Analyses in the separate HC and PD groups

Whole-brain regressions testing for an association between loneliness and looming activation of *a priori* subcortical areas (see Methods) were then conducted (*p* < 0.05, FDR-corrected; also see exploratory results at *p* < .01 cluster-wise corrected threshold in Table S4 and Figure S4). In the HC group, a significant cluster was observed within the right thalamus (peak *Z* = -3.830, *p* = 1 × 10^−4^); this association arose from a negative correlation between Approach > Withdrawal responses of the right thalamus and loneliness (Table 2; Figure 2). In the PD group, a similar (and stronger) pattern of results was found; levels of loneliness were significantly negatively correlated with Approach > Withdrawal activation, most strongly in the right hippocampus (peak *Z* = -4.345, *p* = 1 × 10^−5^), as well as in the right putamen, right caudate nucleus and left pallidum (Table 2; Figure 2). This pattern of results can also be represented as positive correlations between loneliness and Withdrawal > Approach responses (Figure 3A).

**Table 2.**
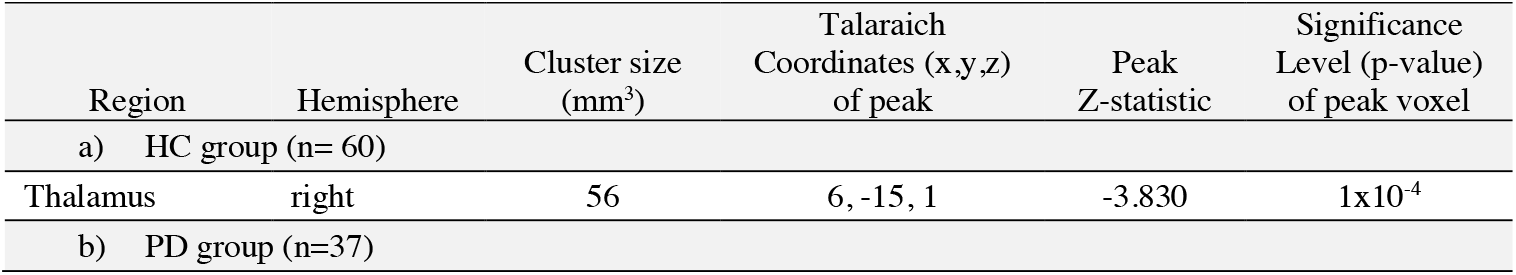
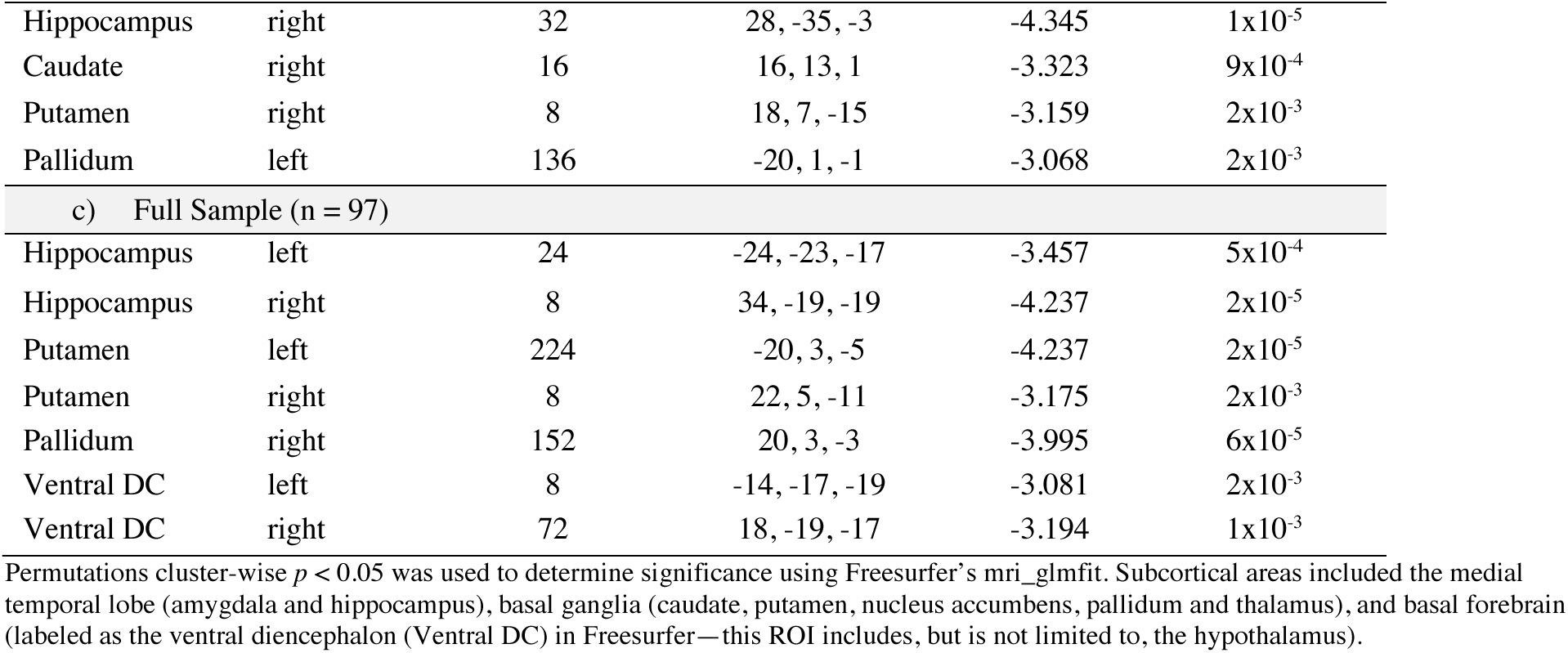
Locations of clusters (surviving whole brain correction) showing negative correlations between responses to Looming (Approach > Withdrawal) faces and loneliness in the subcortical regions-of-interest in the Healthy Control (HC), Psychotic Disorder (PD) and full samples.

| Region | Hemisphere | Cluster size<br>(mm<sup>3</sup>) | Talairach<br>Coordinates (x,y,z)<br>of peak | Peak<br>Z-statistic | Significance<br>Level (p-value)<br>of peak voxel |
| --- | --- | --- | --- | --- | --- |
| a) HC group (n= 60) |  |  |  |  |  |
| Thalamus | right | 56 | 6, -15, 1 | -3.830 | 1x10<sup>-4</sup> |
| b) PD group (n=37) |  |  |  |  |  |
| Hippocampus | right | 32 | 28, -35, -3 | -4.345 | 1x10<sup>-5</sup> |
| Caudate | right | 16 | 16, 13, 1 | -3.323 | 9x10<sup>-4</sup> |
| Putamen | right | 8 | 18, 7, -15 | -3.159 | 2x10<sup>-3</sup> |
| Pallidum | left | 136 | -20, 1, -1 | -3.068 | 2x10<sup>-3</sup> |
| c) Full Sample (n = 97) |  |  |  |  |  |
| Hippocampus | left | 24 | -24, -23, -17 | -3.457 | 5x10<sup>-4</sup> |
| Hippocampus | right | 8 | 34, -19, -19 | -4.237 | 2x10<sup>-5</sup> |
| Putamen | left | 224 | -20, 3, -5 | -4.237 | 2x10<sup>-5</sup> |
| Putamen | right | 8 | 22, 5, -11 | -3.175 | 2x10<sup>-3</sup> |
| Pallidum | right | 152 | 20, 3, -3 | -3.995 | 6x10<sup>-5</sup> |
| Ventral DC | left | 8 | -14, -17, -19 | -3.081 | 2x10<sup>-3</sup> |
| Ventral DC | right | 72 | 18, -19, -17 | -3.194 | 1x10<sup>-3</sup> |

Permutations cluster-wise $p < 0.05$ was used to determine significance using Freesurfer's mri\_glmfit. Subcortical areas included the medial temporal lobe (amygdala and hippocampus), basal ganglia (caudate, putamen, nucleus accumbens, pallidum and thalamus), and basal forebrain (labeled as the ventral diencephalon (Ventral DC) in Freesurfer—this ROI includes, but is not limited to, the hypothalamus).

**Figure 2.**
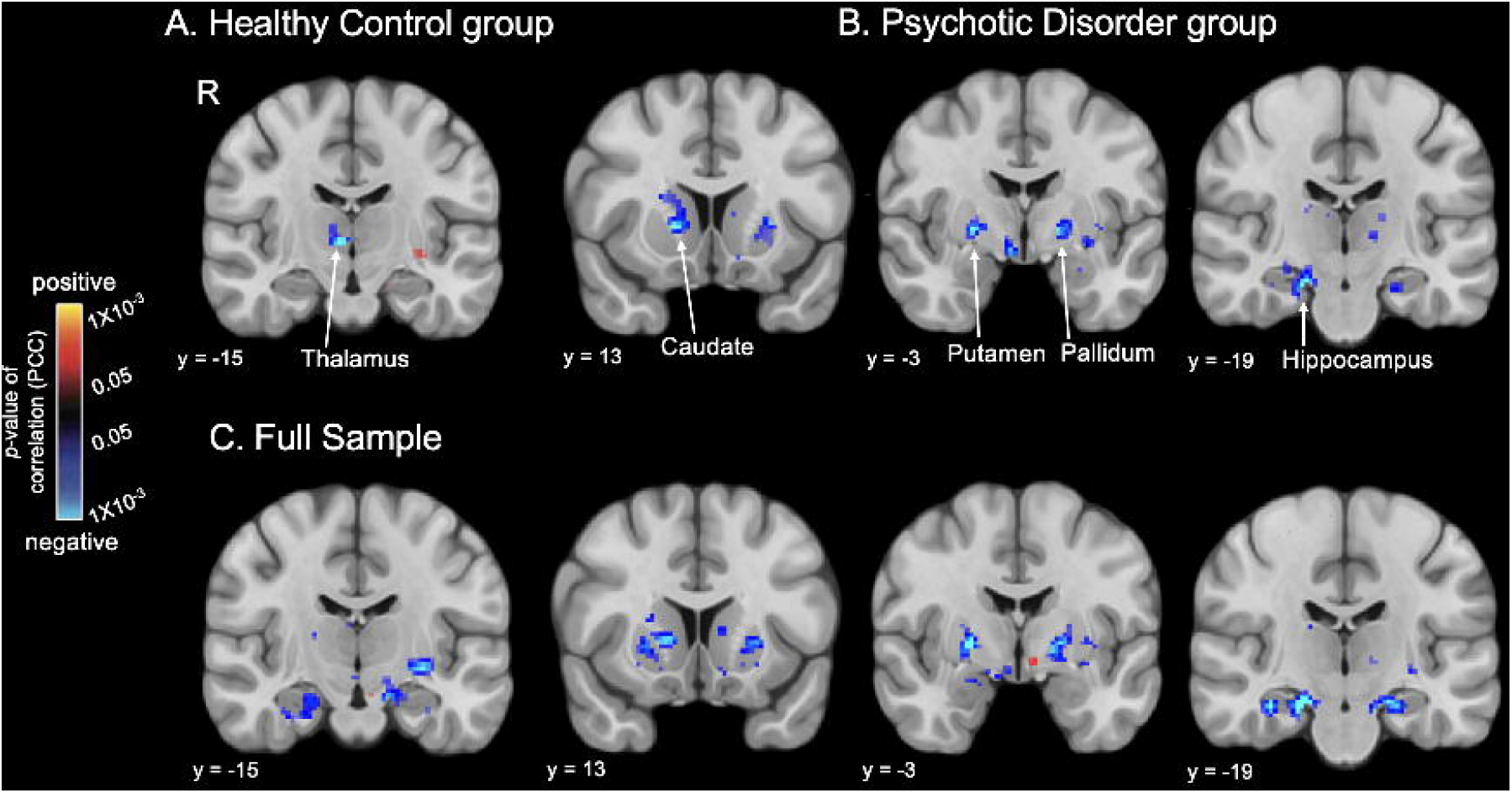
Associations between looming responses of subcortical areas and loneliness. Maps of the results of whole brain regressions reveal significant associations between self-reported loneliness and looming (Approach > Withdrawal) BOLD activation in subcortical areas, overlaid on a MNI common space brain template, (display threshold: *p* < 0.05, uncorrected) in the healthy control (A, n = 60), psychotic disorder (B, n = 37) groups and full sample (C, n = 97). Warm colors (red, yellow) indicate voxels showing significant positive correlations, and cool colors (blue) indicate voxels showing significant negative correlations between Approach > Withdrawal activation and loneliness. Thus, voxels labeled blue also represent those showing significant positive correlations between loneliness and Withdrawal > Approach activation. R, right hemisphere; PCC, partial correlation coefficient.

**Figure 3.**
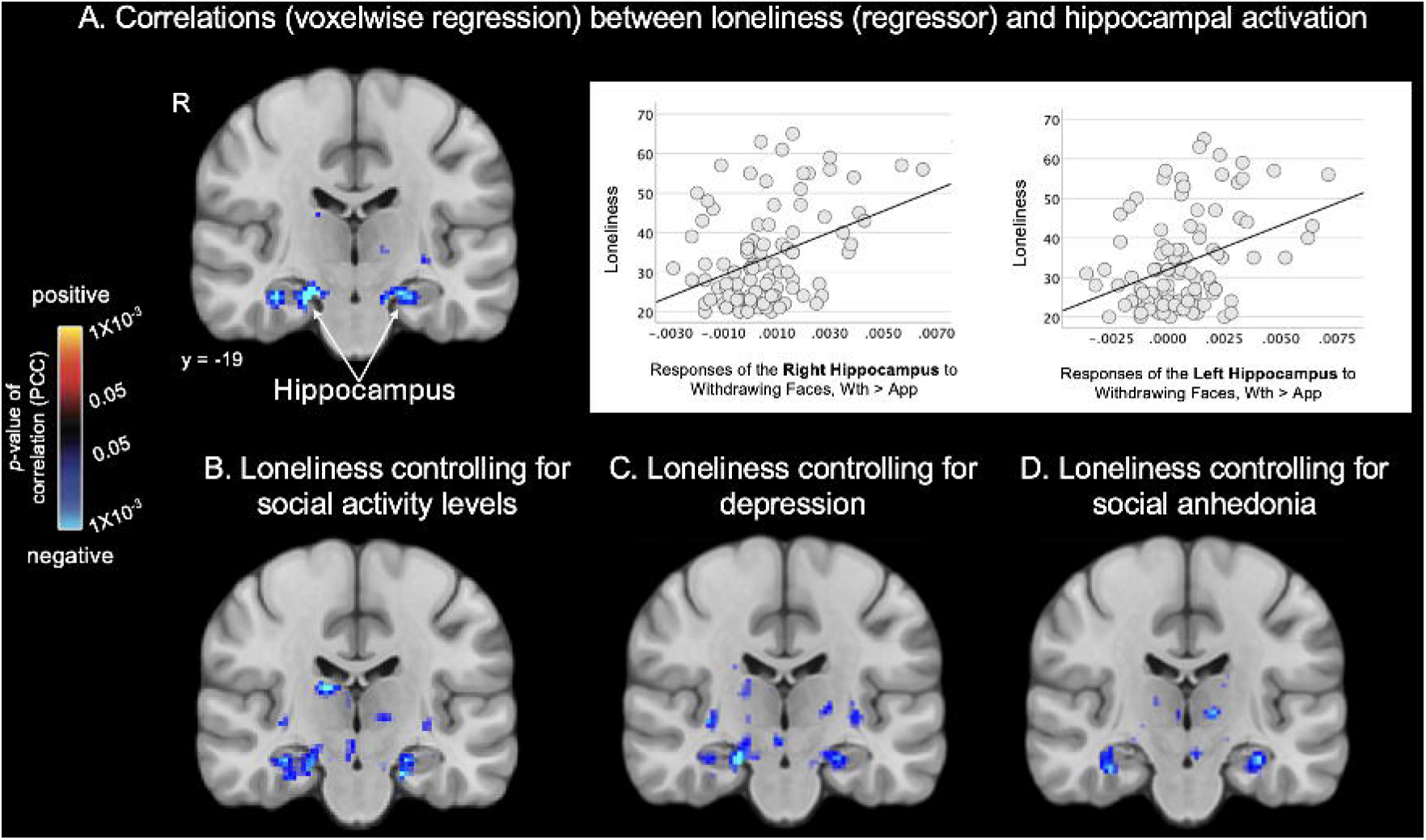
Loneliness is associated with looming responses of the hippocampus, even after controlling for potential confounds. Maps of the results of whole brain regressions conducted in the full sample (n = 97) reveal significant associations between loneliness and looming (Approach > Withdrawal) activation in the hippocampus. Subcortical significance maps (display threshold: *p* < 0.05, uncorrected) resulting from the voxel-wise regression analysis testing for associations between loneliness and looming activation without covariates (A) and when controlling for social activity level (B), depression (C) or social anhedonia (D) are shown. Plots in (A) demonstrate the significant association between Withdrawing > Approaching faces and loneliness in the whole group. Warm colors (red, yellow) indicate voxels showing significant positive correlations (none are shown here), and cool colors (blue) indicate voxels showing significant negative correlations between Approach > Withdrawal activation and loneliness. Thus, voxels labeled blue also represent those showing significant positive correlations between loneliness and Withdrawal > Approach activation. R, right hemisphere; PCC, partial correlation coefficient; Wth, withdrawal; App, approach.

#### Analyses in the full sample

Given that there were no significant differences between the HC and PD groups in Looming activation at a whole brain-corrected statistical threshold, the above regression analyses were repeated in the full sample (n = 97) in order to gain additional power. In the full sample, loneliness was most strongly correlated with Looming-related activation of the right hippocampus (peak *Z* = -4.237, *p* = 2 × 10^−5^). Similar effects were also observed in the left hippocampus, bilateral basal forebrain, right pallidum, and bilateral putamen (Table 2 and Figure 2).

#### Exploratory analyses in cortical regions

There were no significant associations between loneliness and Looming activation of the cortical ROIs in the HC, PD or full samples (*p* < 0.05 cluster-wise FDR-corrected).

#### Controlling for potential confounds

Because loneliness was correlated to varying degrees with social isolation (i.e., low social activity levels), depression and social anhedonia (Table S1), the regression analysis testing for associations between loneliness and looming activation was repeated in the full sample using these variables as covariates. The associations between loneliness and responses of the right and left hippocampus were the only findings that remained significant when controlling for each of these variables (Table S5 and Figure 3).

#### Baseline neural predictors of change in loneliness over one year

The longitudinal regression analysis in the subsample with both baseline and one-year follow-up data (35 HC and 31 PD participants) revealed that Looming-related activation of only the right hippocampus predicted greater loneliness at one year follow-up (peak *Z* = -3.223, *p* = 2 × 10^−4^; Figure 4).

**Figure 4.**
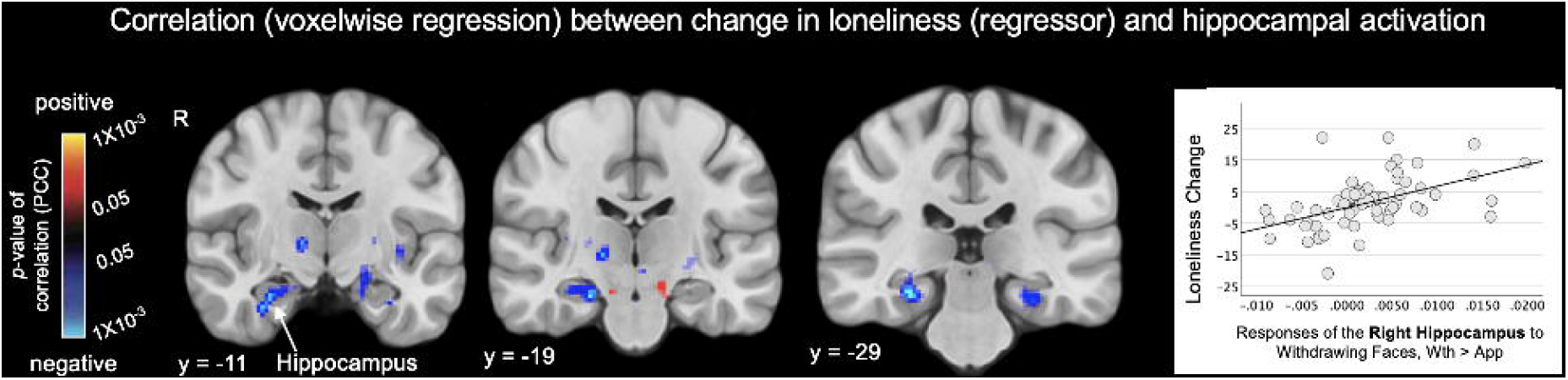
Responses of the hippocampus at baseline predict change in loneliness one year later. Maps of the results of whole brain voxel-wise regressions (display threshold: *p* < 0.05, uncorrected) conducted in a subset of the full sample (n = 66) reveal the significant associations between Approach > Withdrawal activation of the right hippocampus (*p* < 0.05, FDR-corrected) at baseline and changes in loneliness between baseline and the one-year follow-up time point. The scatter plot illustrates this association, displaying the positive correlation between the responses of the right hippocampus to Withdrawing > Approaching faces at baseline and changes in loneliness over one year (larger responses were associated with increases in loneliness). In A, warm colors (red, yellow) indicate voxels showing significant positive correlations, and cool colors (blue) indicate voxels showing significant negative correlations between Approach > Withdrawal activation and change (One Year > Baseline) in loneliness. Thus, voxels labeled blue also represent those showing significant positive correlations between an increase in loneliness over one year and Withdrawal > Approach activation. R, right hemisphere; PCC, partial correlation coefficient; Wth, withdrawal; App, approach.

## DISCUSSION

### Summary of main findings

This study aimed to identify brain mechanisms specifically linked to loneliness in both healthy individuals and those diagnosed with a psychotic disorder. In the psychotic disorder group and the full sample (and at a lower threshold in the healthy control group), loneliness was correlated with responses of the hippocampus to withdrawing face stimuli. In addition to the hippocampus, responses of basal ganglia regions (the right caudate and putamen and left pallidum in the psychotic disorder group, the right thalamus in the healthy control group) to withdrawing faces showed a similar relationship to loneliness. Critically, only the association between loneliness and responses of the hippocampus remained significant in the full sample after controlling for potential confounding factors. The central role of the hippocampus in the experience of loneliness was further highlighted by the finding in a subset of the sample that these hippocampal responses also predicted increases in loneliness one year later.

### Prior evidence for a role of the hippocampus in loneliness

The hippocampus has been consistently implicated in prior neuroimaging studies of loneliness conducted in a range of populations (37). Several studies, including two conducted in large samples (N = ∼2,000 (21) and ∼40,000 (18)), have detected significant negative correlations between loneliness and hippocampal volumes (18–21,23). Also, loneliness has been associated with reductions in the integrity of white matter fibers both intrinsic to and originating from the hippocampus (22,38).

In addition to these anatomical findings, loneliness has also been linked to altered functional connectivity and task-elicited responses of the hippocampus (6,13,16,17,20). For example, altered functional connectivity of the CA1 subfield and molecular layer of the hippocampus with regions of the default network has been demonstrated in individuals endorsing loneliness in the UK Biobank sample (13). In addition, diminished hippocampal-nucleus accumbens connectivity during negative social feedback was observed in a group of healthy individuals with high levels of loneliness (16). Taken together, prior research and the current study indicate that changes in hippocampal structure and function are associated with loneliness.

The neurobiological mechanisms underlying these associations and the direction of these effects remain unknown. Given that loneliness has been linked to increases in cortisol and peripheral physiological measures of stress in humans (39), a reduction in hippocampal volume in lonely individuals may reflect changes in hippocampal morphology (18–21,23) and neurogenesis (24) that are similar to those previously detected in animal models of the effects of chronic stress on the brain. Future longitudinal studies can further investigate whether loneliness-related stress responses lead to changes in hippocampal structure and function in humans and determine whether loneliness precedes or follows such changes.

### The basal ganglia in loneliness

The current study also found that responses of basal ganglia regions, the caudate nucleus, putamen and thalamus, to withdrawing faces correlated with loneliness. However, these effects did not remain significant after controlling for potentially confounding factors, such as isolation, social anhedonia and depression. Thus, one possible interpretation of this pattern of results is that the basal ganglia play a role in experiences or symptoms that often co-occur with loneliness. Consistent with this, one prior study showed that decreased gray matter volumes of the pallidum, putamen and caudate nucleus were significantly correlated with loneliness in a group of older adults with a history of multiple depressive episodes (40). Given the known role of the basal ganglia in reward processes (41), it is possible that basal ganglia circuitry plays a role in signaling the discrepancy between the expected and actual level of social connection experienced by an individual and trigger responses (including depressed mood and its physiological correlates) and actions that may lead to correction of this discrepancy (24,42).

### Associations between loneliness and hippocampal responses in psychotic disorders

Many prior neuroimaging studies of schizophrenia that have employed a case-control design and a hypothesis-driven, region-of-interest approach have detected significant reductions (on average) in hippocampal volume or abnormalities in hippocampal responses in groups of individuals diagnosed with schizophrenia, when compared to healthy control subjects (36,43,44). In the current study, when we used a liberal statistical threshold (comparable to many prior hypothesis-driven, region-of-interest focused studies), we detected significant between-group differences in activation of the hippocampus and other regions, with significantly greater responses of those regions to approaching versus withdrawing faces in the healthy control group compared to the psychotic disorder group. These between-group differences in the responses of the hippocampus and other regions were driven by the greater responses to withdrawing (compared to approaching) face stimuli in the psychotic disorder group (which were correlated with loneliness in the current study), suggesting that the high levels of loneliness in this group may contribute to these findings. This possibility was supported by secondary analyses that included loneliness or social activity levels as a covariate, which showed that the inclusion of these covariates eliminated the between-group differences of the original analyses. Thus, this pattern of results suggests that the effects on the brain of chronic loneliness and/or social isolation, experienced by ∼ 80% of people with serious mental illnesses (4), may be a critical variable to consider when interpreting prior findings and conducting future investigations of the neurobiological basis of schizophrenia.

### Longitudinal analysis

A longitudinal analysis in a subset of the participants from both groups revealed that the hippocampal responses to withdrawing faces at baseline also predicted worsening loneliness over time. This finding preliminarily suggests that this hippocampal response represents a marker of the neurocognitive processes that drive loneliness and possibly of risk for worsening loneliness. Additional studies will be necessary to further confirm this finding and identify the functions of the hippocampus (e.g., autobiographical memory processes(45), mapping of social relationships(46)) that may play a role in the experience of loneliness. Also, future studies could test whether behavioral interventions targeting loneliness can modify this hippocampal response over time.

### Limitations

This study has several limitations to consider when interpreting its results. First, definitive causal inferences cannot be made regarding the relationships between the pattern of neural responses observed and loneliness. Second, a self-report scale was used to measure loneliness; future studies could also include *in vivo* measures (such as ecological momentary assessments or wearable devices measuring aspects of social activity) to capture different determinants of loneliness. However, the self-report measure used here, the UCLA loneliness scale, has been well-validated across many populations and found to correlate with momentary measures of loneliness (47,48), suggesting that it roughly captures the day-to-day conscious experience of feeling lonely for many.

### Future directions

Follow-up studies can be conducted which identify the psychological and biological processes and mediators (e.g., stress-related, inflammatory, hormonal) that may account for the associations observed in this study. An understanding of such mechanisms could lead to the development of interventions that specifically target these processes, potentially improving the physical health and overall quality of life of lonely individuals with and without serious mental illnesses.

## Supporting information

All Supplemental Information

## ACKNOWLEDGMENTS

This research was funded by National Institutes of Mental Health Grant R01MH125426 (DJH), and supported by grants 1S10RR023043, 1S10RR023401, and P41EB015896 for shared resources and imaging resources provided by the Athinoula A. Martinos Center for Biomedical Imaging.

