## Supplementary material for "Association between loneliness and hippocampal responses to dynamic social stimuli in psychotic disorders": All Supplemental Information

### 1. Supplementary Methods

#### *Inclusion/exclusion criteria.*

All participants were 18–50 years of age, proficient in English and were without a history of lifetime drug or alcohol dependence, drug or alcohol abuse within the past six months, current unstable medical illnesses, current or past neurological illnesses, a history of seizures, stroke, or head injury resulting in prolonged loss of consciousness or any standard MRI contraindications (e.g., metal in the body, claustrophobia). Also, all subjects were required to have normal or corrected-to-normal vision. In addition, potential healthy control (HC) participants were excluded if they had a history of psychotic disorder, or a first- or second-degree relative with a history of a psychotic disorder. All psychotic disorder (PD) participants had a history of having one or more psychotic episodes and had no clinically significant changes in symptoms for at least four weeks prior to enrollment in the study. Diagnoses were initially ascertained using medical records and then confirmed using the Mini International Neuropsychiatric Interview (MINI) (1). Potential PD participants were excluded if they were determined to have a PD diagnosis due to another medical condition or a substance and/or medication.

#### *Personal space measurement.*

The Stop Distance Procedure (SDP) is a highly validated, reliable (kappa ~.7) procedure commonly used to measure interpersonal distance ("personal space") preferences. The SDP is conducted as follows: In a laboratory setting, a confederate who is unknown to the subject stands 3 meters away from the subject, facing the subject. The confederate then slowly walks directly towards the subject, while maintaining a neutral facial expression and eye contact with the subject. Before the beginning of the procedure, the subject was instructed to stop the confederate at two points: first when the subject feels "slightly uncomfortable", i.e., when their personal space boundary has just been reached ("the distance at which you would normally have a conversation with a person you have just met"); second when the subject feels "very uncomfortable", i.e., when their personal space has been entered. A member of the research staff then measures both stopping points as the distance between the end of the shoes of the confederate and the subject. The first stop distance (Distance 1, D1) represents personal space size. The second distance (Distance 2, D2) is used to measure the "permeability" of the personal space boundary. Throughout the SDP, the subject is asked to maintain eye contact with the confederate. Each pair of distances is measured 6 times with each confederate, and the full procedure is carried out twice, with a male and female confederate, the order of which is counter-balanced across subjects.

***MRI data acquisition***

All MRI data were acquired on a 3T Siemens Prisma scanner, using a 64-channel head coil. Whole-brain high-resolution T1-weighted anatomical data were acquired using a single T1-weighted 3D multi-echo MPRAGE scan sequence (1mm$^3$; field-of-view=256x256x176mm; flip angle=7°; TR=2530ms; TE=1.69ms) (2). Task-elicited fMRI data (collected during presentation of the Looming paradigm) were acquired using a whole-brain functional T2*-weighted in-plane, simultaneous, multi-slice (SMS) imaging sequence (2.5mm$^3$; field-of-view=230x230x142.5mm; flip angle=65°; TR=1600ms; TE=30ms; multi-band factor: 3) (3).

***Structural MRI data processing.*** The high-resolution whole-brain T1-weighted anatomical data were processed using the Freesurfer 'recon-all' pipeline, which generates gray-white matter segmentations and cortical surface and subcortical models based on subject-specific anatomy(4). Through this pipeline, automatic labels of boundaries of brain regions were registered to each subject's subcortical space to define subcortical regions-of-interest (5).

***Data quality assurance.*** Tissue segmentation, surface-based registration, and registration from the functional to the anatomical space were visually inspected, and if necessary, corrected using alternate initialization steps. In-scanner head motion was evaluated using both the *fslmotionoutliers* tool and by assessing mean motion correction vector translation (MCVT) (6). No participant's entire dataset was identified as meeting criteria for exclusion from the analyses using the following criteria: 1) $\geq$ 20% outliers, and 2) mean MCVT > 1mm. Isolated task runs with excessive head motion within individual datasets were excluded from the analyses: 1 run for 7 participants (2 HC, 5 PD), 2 runs for 6 participants (2 HC; 4 PD), and 3 runs for 1 PD participant.

***Functional MRI data preprocessing.*** The whole-brain Looming task functional MRI data were preprocessed using Freesurfer's Functional Analysis Stream (FS-FAST) preprocessing steps (FreeSurfer v7.4.1), which implemented motion-correction, slice timing correction, smoothing using a 5mm FWHM kernel, and boundary-based registration of the BOLD time-series data to subject-specific space (4).

***Additional details about the Looming paradigm***

During the Looming paradigm, the width of the faces presented varied between 1.1° (64 pixels) and 37.4° (2186 pixels) of visual angle (screen viewing distance = 104 cm; pixels-per-degree = 58.5) over the course of each trial, with the rate of change equivalent to a typical walking speed (112cm/s; 2.5 mph). The face images were generated using the FaceGen software package (http://www.facegen.com) and were presented using MATLAB (version 2018a) and the

Psychophysics MATLAB toolbox, via back-projection onto a screen within the MRI scanner bore (7).

#### *Region-of-interest (ROI) definition*

A previously published fMRI dataset (8–11), which was collected using the same fMRI task (the Looming task) from a non-clinical sample of young adults ($n$ = 130), was used to independently define the ROIs of the network of brain regions previously shown to be involved in personal space regulation (9,10,12). Using the previous data, a group-level GLM analysis contrasting the Approaching vs. Withdrawing face conditions was thresholded at a significance level of $p < 0.0001$ and subjected to cluster correction using permutation testing (cluster-wise $p < 0.05$; 1000 permutation trials; see Table S2 for complete list of the significant clusters). These cluster-corrected significance maps revealed two clusters of activation in the parietal cortex located within superior and inferior parietal cortex, respectively, oriented in the anterior-posterior plane along the intra-parietal sulcus (Figure S2; ROIs outlined in green). ROIs within the premotor cortex, which is an established node of the personal space network based on prior studies conducted in humans (9,10,12) and non-human primates (13,14), were defined using resting-state fMRI data also acquired as part of the same previously collected dataset (9,10,12).

### 2. **Supplementary Tables**

**Table S1.** Correlations (r/p values) among the symptom and behavioral measures.

| Measure | 1 | 2 | 3 | 4 | 5 | 6 |
|---|---|---|---|---|---|---|
| a) Healthy Controls (n = 60) | | | | | | |
| 1. Loneliness[a] | ___ | | | | | |
| 2. Social Activity Levels[b] | -0.231/0.071 | ___ | | | | |
| 3. Depression[c] | 0.698/< 0.001* | -0.160/0.222 | ___ | | | |

|  |  |  |  |  |  |  |
|---|---|---|---|---|---|---|
| 4. Social Anhedonia[d] | 0.692/<0.001* | -0.235/0.071 | 0.422/<0.001* | ___ |  |  |
| 5. Trustworthy Ratings[e] | -0.026/0.841 | 0.066/0.606 | -0.253/0.045* | -0.047/0.712 | ___ |  |
| 6. Personal Space Size[f] | 0.172/0.096 | 0.039/0.706 | 0.156/0.131 | 0.194/0.137 | -0.143/0.276 | ___ |
| 7. Loneliness Change[g] | -0.188/0.266 | 0.049/0.774 | -0.030/0.858 | -0.089/0.599 | 0.014/0.933 | -0.193/0.259 |
| b) Psychotic Disorders (n = 37) |  |  |  |  |  |  |
| 1. Loneliness[a] | ___ |  |  |  |  |  |
| 2. Social Activity Levels[b] | -0.242/0.143 | ___ |  |  |  |  |
| 3. Depression[c] | 0.668/< 0.001* | -0.071/0.684 | ___ |  |  |  |
| 4. Social Anhedonia[d] | 0.513/.001* | -0.373/0.027* | 0.470/0.004* | ___ |  |  |
| 5. Trustworthy Rating[e] | 0.011/0.934 | 0.096/0.457 | -0.148/0.381 | -0.191/0.258 | ___ |  |
| 6. Personal Space Size[f] | -0.259/0.132 | 0.318/0.063 | 0.017/0.921 | -0.188/0.279 | -0.156/0.370 | ___ |
| 7. Loneliness Change[g] | -0.570/0.003* | 0.113/0.592 | -0.206/0.324 | 0.148/0.480 | -0.081/0.700 | 0.012/0.955 |
| c) Full Sample (n = 97) |  |  |  |  |  |  |
| 1. Loneliness[a] | ___ |  |  |  |  |  |
| 2. Social Activity Levels[b] | -0.323/0.001* | ___ |  |  |  |  |
| 3. Depression[c] | 0.712/<0.001* | -0.207/0.044* | ___ |  |  |  |
| 4. Social Anhedonia[d] | 0.630/<0.001* | -0.328/0.001* | 0.468/<.001* | ___ |  |  |
| 5. Trustworthy Rating[e] | -0.185/0.066 | 0.115/0.256 | -0.254/0.011* | -0.131/0.193 | ___ |  |
| 6. Personal Space Size[f] | 0.253/0.045* | 0.014/0.915 | 0.141/0.282 | 0.194/0.137 | -0.197/0.055 | ___ |
| 7. Loneliness Change[g] | -0.375/0.003* | 0.098/0.451 | -0.148/.251 | .018/.891 | -0.007/0.956 | -0.098/.456 |

Pearson's correlations (the r and p-values are listed for each) in the a) Healthy Control (HC) group, b) Psychotic Disorders (PD) group and c) Full Sample. The variables of interest were measured using: [a]the UCLA Loneliness scale, [b]the Social Network Index: number of social contacts, [c]the Beck Depression Inventory (BDI), [d]the Social Anhedonia Scale-Revised (SAS), [e]the trustworthy ratings of the Looming task face stimuli, and [f]the Stop Distance Procedure, [g]change in the UCLA Loneliness scale from baseline to one year. Significant correlations ($p < 0.05$) are indicated with an asterisk (*).

**Table S2.** Significant clusters within cortical and subcortical regions-of-interest showing looming (Approach vs. Withdrawal) activation in the healthy control group, psychotic disorder group and the full sample.

| | Hemisphere | Size ($mm^3$) | Talaraich Coordinates (x,y,z) | Z statistic | Significance Level (p-value) |
|---|---|---|---|---|---|
| a) Healthy Control group |  |  |  |  |  |
| Dorsal premotor | left | 372.37 | -46.1, -2.4, 35.2 | 8.479 | $1x10^{-15}$ |
| Dorsal premotor | left | 200.98 | -19.3, -2.8, 58.3 | 6.153 | $8x10^{-10}$ |
| Dorsal premotor | right | 1804.48 | 30.9, -5.5, 44.4 | 10.934 | $<1x10^{-15}$ |
| Inferior parietal | left | 449.09 | -23.7, -60.7, 51.1 | 10.161 | $<1x10^{-15}$ |
| Inferior parietal | right | 611.30 | 23.7, 74.6, 26.0 | 11.562 | $<1x10^{-15}$ |
| Superior parietal | right | 127.49 | 23.3, -57.2, 50.3 | 10.603 | $<1x10^{-15}$ |
| Ventral premotor | left | 444.81 | -40.1, -7.5, 48.5 | 10.577 | $<1x10^{-15}$ |
| Ventral premotor | right | 100.55 | 15.3, -26.4, 44.7 | 5.942 | $3x10^{-9}$ |
| Caudate | left | 32 | -20, 11, 13 | 3.463 | $5x10^{-4}$ |

| | | | | | |
|---|---|---|---|---|---|
| Putamen | left | 64 | -24, 3, -5 | 3.446 | $5x10^{-4}$ |
| Hippocampus | left | 328 | -26, -19, -13 | -4.959 | $7x10^{-7}$ |
| Ventral DC | left | 104 | -22, -29, -3 | 4.560 | $5x10^{-6}$ |
| Ventral DC | right | 1784 | 28, -21, -11 | -6.619 | $3x10^{-11}$ |
| Ventral DC | right | 120 | 4, -15, -7 | 5.022 | $5x10^{-7}$ |
| Thalamus | right | 328 | 16, -31, 3 | 5.732 | $1x10^{-9}$ |
| Thalamus | right | 8 | 14, -7, 7 | 3.174 | $1x10^{-3}$ |
| Thalamus | right | 16 | 8, -11, -3 | 3.111 | $1x10^{-3}$ |
| b) Psychotic Disorder group | | | | | |
| Dorsal premotor | left | 137.14 | -34.7, -1.3, 45 | 4.290 | $2x10^{-5}$ |
| Dorsal premotor | right | 245.36 | 34.6, -6.4, 45 | 5.246 | $2x10^{-7}$ |
| Inferior parietal | left | 308.27 | -22.5, -77, 29.6 | 5.755 | $9x10^{-9}$ |
| Inferior parietal | right | 108.87 | 25.5, -75.4, 33.4 | 4.745 | $2x10^{-6}$ |
| Superior parietal | left | 343.32 | -23.9, 59.6, 51.0 | 7.360 | $2x10^{-13}$ |
| Ventral premotor | right | 758.97 | 40.6, 0.4, 34.3 | 4.843 | $1x10^{-6}$ |
| Amygdala | left | 16 | -26, -11, -21 | -3.564 | $4x10^{-4}$ |
| Hippocampus | left | 728 | -32, -37, -7 | -5.686 | $1x10^{-8}$ |
| Hippocampus | left | 248 | -32, -15, -17 | -3.785 | $2x10^{-4}$ |
| Ventral DC | left | 80 | -12, -17, -21 | -3.682 | $2x10^{-4}$ |
| Thalamus | left | 16 | -18, -33, 1 | 3.434 | $6x10^{-4}$ |
| Thalamus | right | 8 | 14, -21, 5 | -3.067 | $2x10^{-3}$ |
| Thalamus | left | 32 | -16, -23, 11 | -3.560 | $4x10^{-4}$ |
| c) Full Sample | | | | | |
| Dorsal Premotor | left | 504.86 | -37.3, -76.3, 27.7 | 15.579 | $<1 x10^{-15}$ |
| Dorsal Premotor | right | 1800.24 | 32.5, -5.0, 43.9 | 14.036 | $<1 x10^{-15}$ |
| Inferior parietal | left | 650.05 | -23.0, -76.0, 27.7 | 15.579 | $<1 x10^{-15}$ |
| Superior parietal | left | 455.11 | -23.8, -59.9, 51.3 | 16.307 | $<1 x10^{-15}$ |
| Ventral Premotor | left | 390.17 | -46.1, -2.5, 35.8 | 11.323 | $<1 x10^{-15}$ |
| Hippocampus | left | 16 | -22, -15, -25 | -3.096 | $2 x10^{-3}$ |
| Ventral DC | left | 240 | -22, -29, -3 | 6.433 | $1 x10^{-10}$ |
| Thalamus | left | 24 | -2, -9, 7 | -3.176 | $1 x10^{-3}$ |
| Thalamus | right | 224 | 20, -31, -1 | 5.792 | $7 x10^{-9}$ |

A permutations cluster-wise $p < 0.05$ significance threshold was used (using mri_glmfit of FreeSurfer) for the *a priori* ROIs. Positive Z statistic values indicate significant Approach > Withdrawal activation, whereas negative Z statistic values indicate significant Withdrawal > Approach activation.

**Table S3.** HC vs PD group differences in looming activation ($p < 0.05$ uncorrected).

| Region | Hemisphere | Talaraich Coordinates (x,y,z) | Peak Z statistic | Significance Level (p-value) |
|---|---|---|---|---|
| Caudate | left | -17, 15, 3 | -3.3 | 0.002 |
| Caudate | left | -18, 19, 24 | 2.5 | 0.012 |
| Caudate | right | 20, 21, 5 | -2.1 | 0.041 |
| Caudate | right | 12, 23, 20 | 2.0 | 0.045 |
| Putamen | left | -31, 1, 5 | 2.3 | 0.021 |
| Putamen | left | -34, 13, 4 | -2.4 | 0.020 |
| Putamen | right | 25, 7, -5 | -2.4 | 0.020 |
| Amygdala | left | -28, 3, -13 | 2.2 | 0.033 |
| Hippocampus | left | -25, -17, -22 | 3.1 | 0.002 |
| Hippocampus | right | 18, -3, -8 | 2.9 | 0.006 |
| Ventral DC | left | -4, -1, 0 | 2.0 | 0.045 |
| Ventral DC | left | -9, 13, 1 | -2.3 | 0.026 |
| Ventral DC | right | 1, -1, 1 | 2.2 | 0.033 |
| Ventral DC | right | 3, 9, -2 | -1.7 | 0.026 |

Positive Z statistic values indicate clusters with HC > PD looming (Approach > Withdrawal) activation (in the amygdala, hippocampus, caudate, putamen and ventral DC), whereas negative Z statistic values indicate clusters with PD > HC looming activation (in the caudate, putamen and ventral DC). DC = diencephalon. All clusters $\geq$ 10 voxels.

**Table S4**. Locations of peak clusters showing negative correlations between responses to Looming (Approach > Withdrawal) faces and loneliness in the subcortical regions-of-interest in the Healthy Control (HC), Psychotic Disorder (PD) and full samples at the $p < 0.01$ cluster-corrected level.

| Region | Hemisphere | Cluster size ($mm^3$) | Talaraich Coordinates (x,y,z) of peak | Peak Z-statistic | Significance Level (p-value) of peak voxel |
|---|---|---|---|---|---|
| a) HC group (n= 60) | | | | | |
| Hippocampus | left | 24 | -33, -17, -19 | -2.7 | $3x10^{-2}$ |
| Hippocampus | right | 34 | 28, -37, -2 | -3.1 | $9 x10^{-4}$ |
| Pallidum | right | 16 | 18, -7, -5 | 2.149 | $1x10^{-1}$ |
| Putamen | left | 200 | -32, -13, -3 | -2.365 | $9x10^{-2}$ |
| Thalamus | right | 56 | 6, -15, 1 | -3.830 | $1x10^{-4}$ |
| b) PD group (n=37) | | | | | |
| Hippocampus | right | 32 | 28, -35, -3 | -4.345 | $1x10^{-5}$ |
| Caudate | right | 16 | 16, 13, 1 | -3.323 | $9x10^{-4}$ |

| | | | | | |
|---|---|---|---|---|---|
| Putamen | right | 8 | 18, 7, -15 | -3.159 | 2x10$^{-3}$ |
| Pallidum | left | 136 | -20, 1, -1 | -3.068 | 2x10$^{-3}$ |
| Pallidum | right | 776 | 20, 1, -1 | -3.415 | 3x10$^{-3}$ |
| Thalamus | right | 128 | 14, -9, 3 | -2.378 | 8x10$^{-2}$ |
| Ventral DC | right | 168 | 6, -3, -11 | -2.506 | 6x10$^{-2}$ |
| c) Full Sample (n = 97) | | | | | |
| Hippocampus | left | 24 | -24, -23, -17 | -3.457 | 5x10$^{-4}$ |
| Hippocampus | right | 8 | 34, -19, -19 | -4.237 | 2x10$^{-5}$ |
| Caudate | left | 648 | -16, 3, 17 | -2.635 | 4x10$^{-3}$ |
| Pallidum | right | 152 | 20, 3, -3 | -3.995 | 6x10$^{-5}$ |
| Putamen | left | 224 | -20, 3, -5 | -4.237 | 2x10$^{-5}$ |
| Putamen | right | 8 | 22, 5, -11 | -3.175 | 2x10$^{-3}$ |
| Thalamus | right | 56 | 6, -15, 1 | -3.830 | 1x10$^{-4}$ |
| Ventral DC | left | 8 | -14, -17, -19 | -3.081 | 2x10$^{-3}$ |
| Ventral DC | right | 72 | 18, -19, -17 | -3.194 | 1x10$^{-3}$ |

Permutations cluster-wise whole-brain corrected at $p < 0.01$ was used to determine significance using Freesurfer's mri_glmfit. FDR correction was not applied. Subcortical areas included the medial temporal lobe (amygdala and hippocampus), basal ganglia (caudate, putamen, nucleus accumbens, pallidum and thalamus), and basal forebrain (labeled as the ventral diencephalon (Ventral DC) in Freesurfer—this ROI includes, but is not limited to, the hypothalamus).

**Table S5.** Locations of clusters showing associations between looming responses in subcortical regions-of-interest and loneliness, controlling for potential confounds, in the full sample (n = 97).

| Region | Hemisphere | Cluster size (mm$^3$) | Talaraich coordinates (x,y,z) of peak | Peak Z statistic | Significance level (p-value) of peak voxel |
|---|---|---|---|---|---|
| a) Controlling for social activity levels | | | | | |
| Caudate | left | 8 | -20, 5, 15 | -3.274 | 2 x10$^{-4}$ |
| Caudate | left | 32 | -12, 7, 9 | -3.240 | 1 x10$^{-3}$ |
| Caudate | left | 48 | -16, 1, 19 | -3.239 | 1 x10$^{-3}$ |
| Caudate | right | 232 | 12, 19, -3 | -4.415 | 8 x10$^{-6}$ |
| Putamen | left | 8 | -22, 9, 5 | -3.061 | 2 x10$^{-3}$ |
| Putamen | left | 792 | -20, 5, -5 | -5.758 | 2 x10$^{-9}$ |
| Putamen | right | 8 | 32, -1, -9 | -3.047 | 2 x10$^{-3}$ |
| Putamen | right | 32 | 24, 17, -3 | -3.205 | 1 x10$^{-3}$ |
| Putamen | right | 40 | 16, 11, -13 | -3.791 | 2 x10$^{-4}$ |
| Putamen | right | 48 | 14, 1, -11 | -3.570 | 4 x10$^{-4}$ |
| Putamen | right | 72 | 34, -7, 1 | -3.792 | 2 x10$^{-4}$ |
| Hippocampus | left | 16 | -20, -17, -23 | -3.545 | 4 x10$^{-4}$ |

| | | | | | |
|---|---|---|---|---|---|
| Hippocampus | left | 184 | -26, -25, -17 | -4.764 | $2 \times 10^{-6}$ |
| Hippocampus | right | 16 | 32, -19, -17 | -3.029 | $2 \times 10^{-3}$ |
| Hippocampus | right | 40 | 20, -17, -21 | -3.695 | $2 \times 10^{-4}$ |
| Thalamus | right | 32 | 14, -17, 15 | -3.681 | $2 \times 10^{-4}$ |
| b) Controlling for depression | | | | | |
| Pallidum | right | 104 | 20, 1, 1 | -3.786 | $2 \times 10^{-4}$ |
| Hippocampus | left | 8 | -22, -17, -9 | -3.070 | $2 \times 10^{-3}$ |
| Hippocampus | right | 16 | 24, -15, -25 | -3.083 | $2 \times 10^{-3}$ |
| c) Controlling for social anhedonia | | | | | |
| Caudate | right | 8 | 16, 15, -7 | -3.074 | $2 \times 10^{-3}$ |
| Putamen | left | 16 | -26, 15, -3 | -3.261 | $2 \times 10^{-4}$ |
| Hippocampus | left | 24 | -16, -39, 1 | -3.349 | $8 \times 10^{-4}$ |
| Hippocampus | right | 8 | 38, -21, -19 | -3.288 | $1 \times 10^{-3}$ |
| Hippocampus | right | 24 | 38, -25, -19 | -4.153 | $3 \times 10^{-5}$ |
| Ventral DC | left | 16 | -12, -3, -11 | -3.445 | $5 \times 10^{-4}$ |
| Ventral DC | right | 16 | 18, -5, -11 | -3.115 | $2 \times 10^{-3}$ |

A permutations cluster-wise FDR corrected $p < 0.05$ was used as the significance threshold. Subcortical areas included medial temporal lobe (amygdala and hippocampus), basal ganglia (caudate nucleus, putamen, pallidum and thalamus), and basal forebrain (i.e., the ventral diencephalon, ventral DC) regions delineated by the Desikan-Killiany atlas. Social activity level was measured using the Social Network Index (number of social contacts), depression using the Beck Depression Inventory and social anhedonia using the Chapman Social Anhedonia Scale-Revised. The negative Z statistic values reflect significant negative correlations between levels of loneliness and magnitude of Approach > Withdrawal activation in the corresponding regions.

### Supplementary Figures

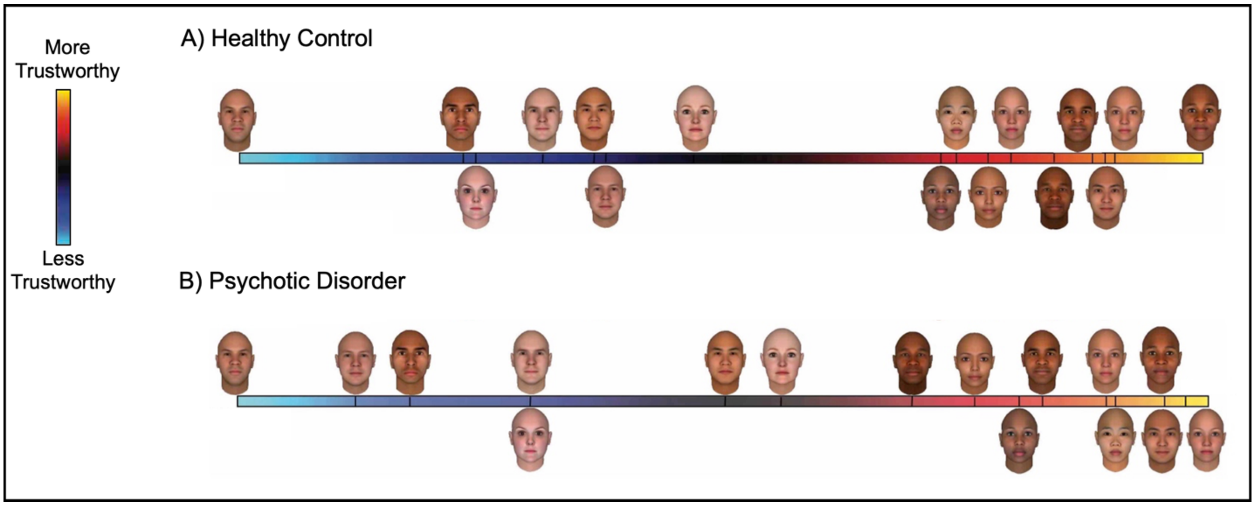


**Figure S1. Trustworthiness face ratings in the healthy control and psychotic disorder groups.** The trustworthiness ratings of the faces presented during the Looming fMRI paradigm in the healthy control (A) and psychotic disorder (B) groups are shown. Ratings were made following the scanning using a scale ranging from 1 to 10, with 1 = least trustworthy and 10 = most trustworthy. These ratings are plotted as untrustworthy (blue) to trustworthy (red), with relative placement based on relative mean rating values for each face in each sample.

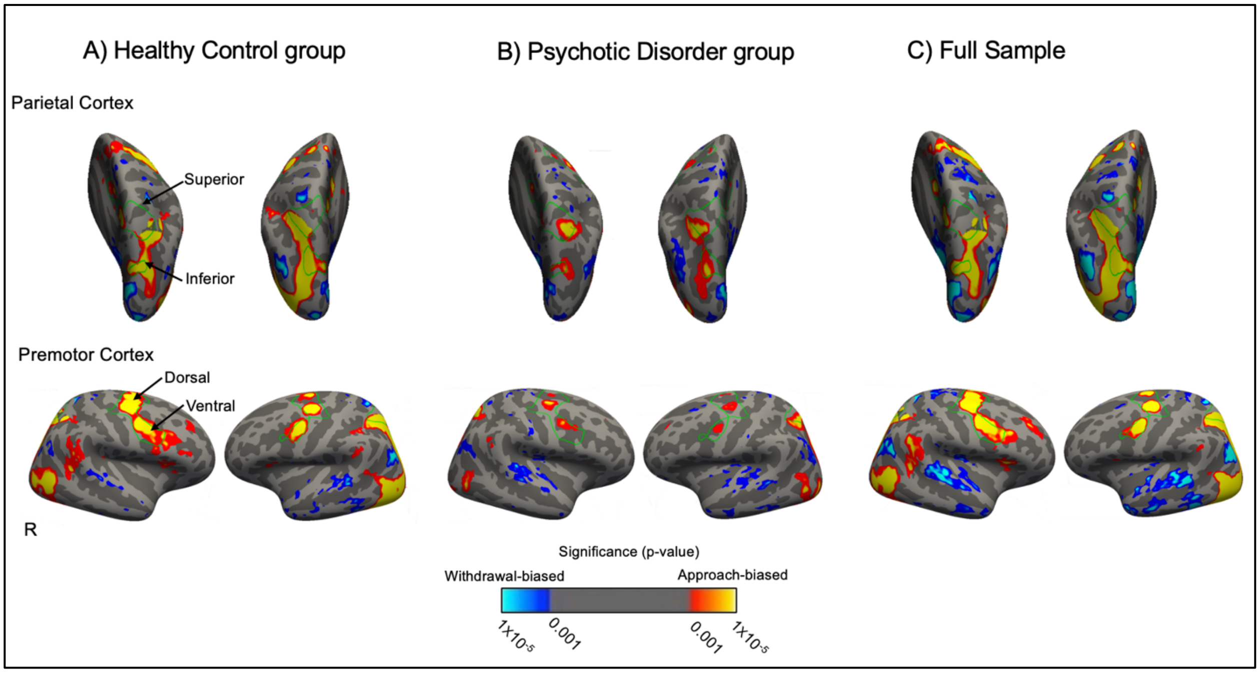


**Figure S2. Looming activation in cortical areas in the healthy control group, psychotic disorder group and full sample.** (A-C) Looming activation is displayed on an inflated cortical surface for the left and right hemispheres. Green outlined regions correspond to each cortical region-of-interest (posterior and lateral views, respectively) defined using an independent fMRI dataset which was collected using the looming paradigm. Significant clusters ($p < 0.001$; corrected) for the Approach > Withdrawal contrast in the A), healthy control group ($n$ = 60), B) psychotic disorder group ($n$ = 37, and C) full sample ($n$ = 97) are displayed. Warm colors (red-yellow) indicate Approach > Withdrawal activation; cool colors (blue) indicate Withdrawal > Approach activation. R, right hemisphere.

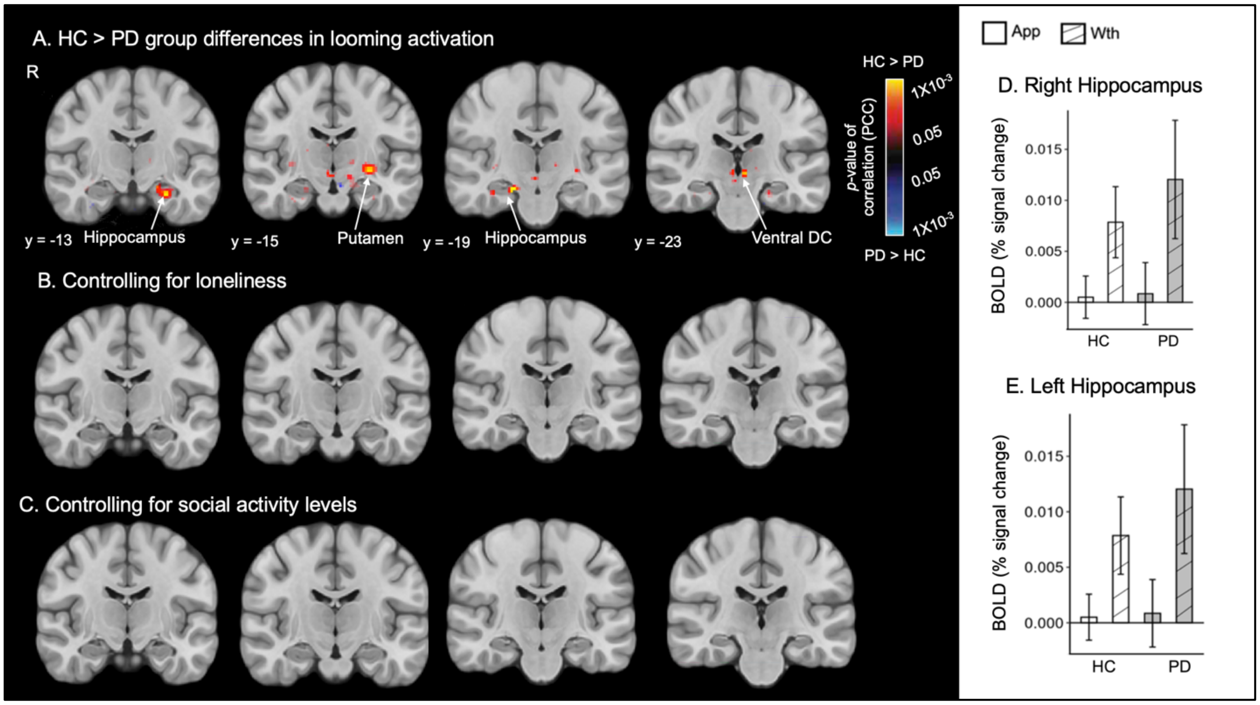


**Figure S3. No HC vs. PD differences in looming activation after controlling for loneliness or social activity levels.** Coronal views of between-group comparison maps ($p < .05$, uncorrected) displaying clusters showing significantly greater Approach > Withdrawal activation in the hippocampus, putamen and ventral diencephalon in the HC compared to the PD group (A), and the same comparison after controlling for loneliness (B) and social activity levels (C). Also, on the right, bar plots are shown of the responses for the two conditions (Approach, Withdrawal) underlying the between-group differences shown in panel A, extracted from a significant cluster in the (D) right hippocampus (peak voxel: 18, -3, -8) and (E) left hippocampus (peak voxel: -25, -17, -22). These plots illustrate that the HC > PD differences in the hippocampus (see panel A) are driven by greater Withdrawal > Approach responses in the PD compared to the HC group (also see Table S3). There were no significant differences between the HC and PD groups in looming activation after controlling for loneliness or social activity. R, right hemisphere; Ventral DC, ventral diencephalon; PCC, partial correlation coefficient; Wth, withdrawal; App, approach.

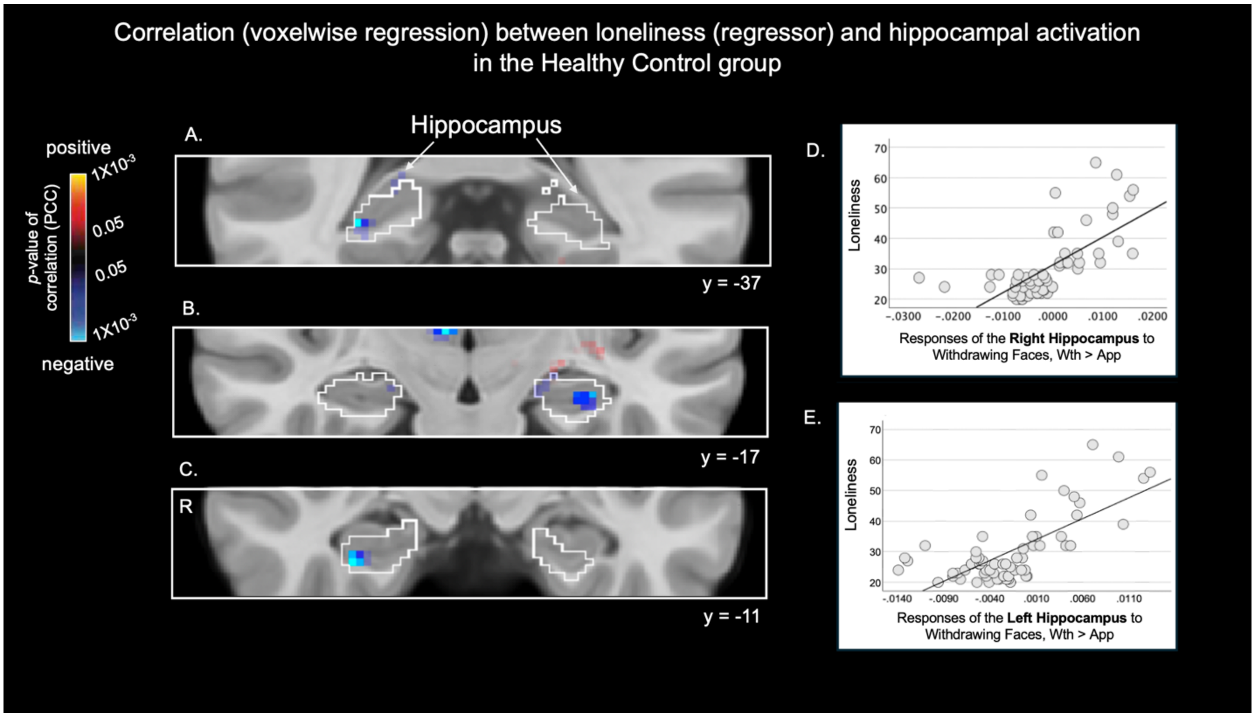


**Figure S4. Loneliness is related to looming activation in the hippocampus in the healthy control group.** Coronal views of regression maps ($p < .05$, uncorrected) of loneliness vs Approach > Withdrawal activation at baseline in the healthy control group (n = 60) overlaid on a MNI common space brain template, displaying three coronal slices (y = -37, -17, -11), showing the hippocampus (outlined in white) (A-C). Cool colors (blue) indicate voxels showing significant negative correlations between Approach > Withdrawal activation and loneliness. Thus, voxels labeled blue also represent those showing significant positive correlations between loneliness and Withdrawal > Approach activation. Plots in D and E illustrate the significant associations between Withdrawal > Approach activation of the right and left hippocampus and loneliness. R, right hemisphere; PCC, partial correlation coefficient; Wth, withdrawal; App, approach.
